# Deconvolution of Sample Identity in Single-Cell RNA Sequencing *via* Genome Imputation

**DOI:** 10.1101/2025.02.11.637700

**Authors:** Rohit Ghosh, Rupert Hugh-White, Farshad Nassiri, Gelareh Zadeh, Paul C. Boutros

## Abstract

**Background:** Droplet based single-cell RNA sequencing (scRNA-seq) is a powerful tool for measuring RNA abundance profiles at cell-specific resolution. Droplet-based barcoding technology allows sample multiplexing, thereby facilitating high-scale of single cell sequencing. The resulting processing complexity, sample contamination and the underlying chemistries can all contribute to cell mis-labelling and consequent spurious cell-to-sample assignment. Approaches for barcode-free de-multiplexing which leverage natural genetic variation have been developed, but generally require an external source of genotype information.

**Results:** We propose a novel method to exploit genome imputation and clustering to assign cells to inferred donor groups in the absence of a priori genetic information. Using tumor-derived single-cell RNA-sequencing (scRNA-seq) data, our workflow successfully assigned individual cells to donor-of-origin with high concordance.

**Conclusions:** This imputation-clustering approach represents a quality-assessment and quality-control strategy for barcode-free single cell donor-origin deconvolution with the capacity to resolve cases of sample cross-contamination.

## Background

Single-cell RNA sequencing (scRNA-seq) provides an estimate of the transcriptome at individual cell resolution in a tissue of interest. In contrast to bulk sample sequencing, single-cell methods are able to resolve cellular heterogeneity related to cell-type and state, and as such are useful for applications including studying cellular transition states or clonal cell populations of a tumor.

Droplet single-cell RNA-sequencing (dscRNA-seq) is a popular form of scRNA-seq protocol and involves the encapsulation of individual cells in oil droplets that are each sequenced. To reduce cost and increase experimental throughput, multiplexing of samples from different individuals is frequently performed. In this approach, droplet-based tagging with oligonucleotide barcodes allows unique labelling of cells from different samples, which are then de-multiplexed during data processing *e*.*g*.(1). However, these approaches add time and cost to sample preparation, which can limit population-scale studies. Further, mis-labelling of reads across samples has been reported in scRNA-seq data produced through barcode-based multiplexing, leading to reduced accuracy of results(2). An alternative approach, as implemented in the tool demuxlet(3), is to use genetic variation as a ‘natural’ barcode to probabilistically assign cells to input samples. However, this method requires prior genotype information of sampled individuals through an independent data source such as bulk DNA sequencing.

We have developed a strategy based on directly clustering cells into donor-level groups using inferred genetic information, without use of barcode or external genetic reference data (**Figure 1**). To ameliorate the effect of genotype-sparsity, we employ haplotype phasing and imputation, with *k*-medoids clustering for efficient handling of high dimensionality single-cell genotype data.

**Figure 1.**
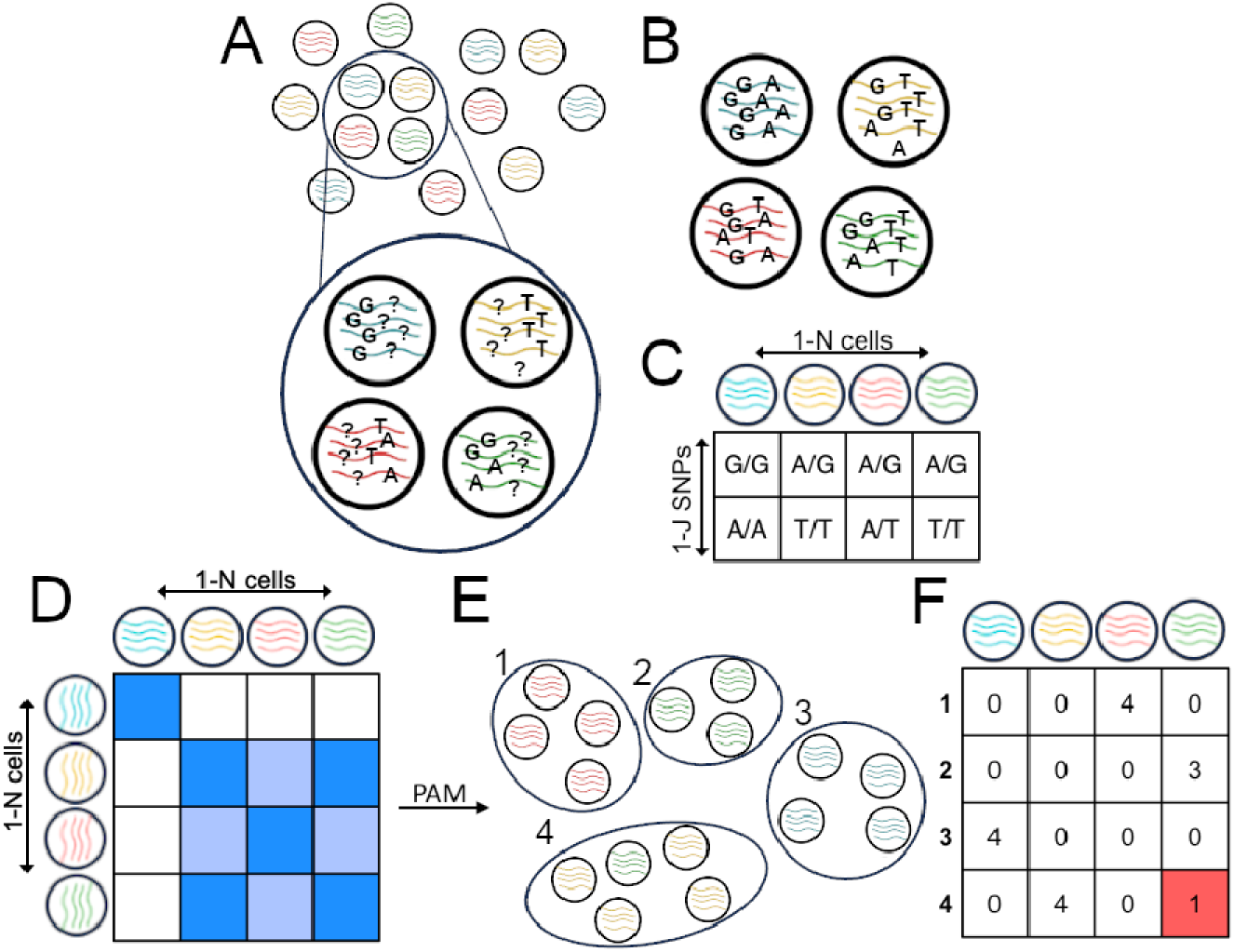
Imputation and Clustering Workflow. **A** Graphical representation of a population of sequenced cells post-genotyping. Different colors of DNA-strands represent different donors of origin inferred through droplet-barcoding. Zoom-in shows cells with partially inferred genotypes. **B** After phasing and imputation, additional single cell genotypes have been inferred. **C** Data are integrated resulting in a cell by SNP genotype matrix. **D** Pairwise hamming distance is calculated to construct a symmetric genotype distance matrix. Intensity of shading represents degree of genetic similarity between each pair of cells. **E** Partitioning around medoids (PAM) is then used to define *k*-clusters of cells corresponding to sample-of-origin identity. In this example four donors were used and thus *k* = 4. As illustrated by cluster 4, in some cases clusters may disagree with barcode identity, potentially reflecting cross-sample contamination. **F** A contingency table represents the agreement between inferred cell cluster (y-axis) and expected barcode-inferred donor (x-axis). Shaded cell indicates a case of a mismatch between assigned cluster and barcode-inferred identity.

## Results

We genotyped a total of 59,880 cells from eight previously published meningioma tumor-derived single nuclei RNA-seq(4) (**Figure 1A**). Genotypes were called at an average of 2,046 SNPs per cell (σ = 1,554), using a reference panel of 7.4 million SNPs identified in the 1,000 Genomes Project(5). Following per-sample phasing and imputation (**Figure 1B**), SNPs were filtered by imputed R^2^ value (> 0.9) and the set of remaining SNPs genotyped in at least one cell per sample (70,981 SNPs) was used for downstream analysis (**Figure 1C**). This genotype matrix was then used to construct a cell distance matrix using Hamming distance (**Figure 1D**). Finally, we employed *k*-medoids clustering to assign each individual cell to a cluster based on genetic similarity (**Figure 1E**) using Partitioning Around Medoids (PAM)(6). PAM was selected to handle the high dimensionality genotype matrix due to its relatively low time complexity.

Here, we assumed *k* to be known a priori and to be equal to the number of sampled individuals (eight). Cell cluster assignments were almost perfectly concordant with expected donor individual of origin (**Figure 2**), indicating that our implemented workflow is successfully able to leverage scRNA-seq derived genotype data to recover cell-sample identity independent of barcoding information.

**Figure 2.**
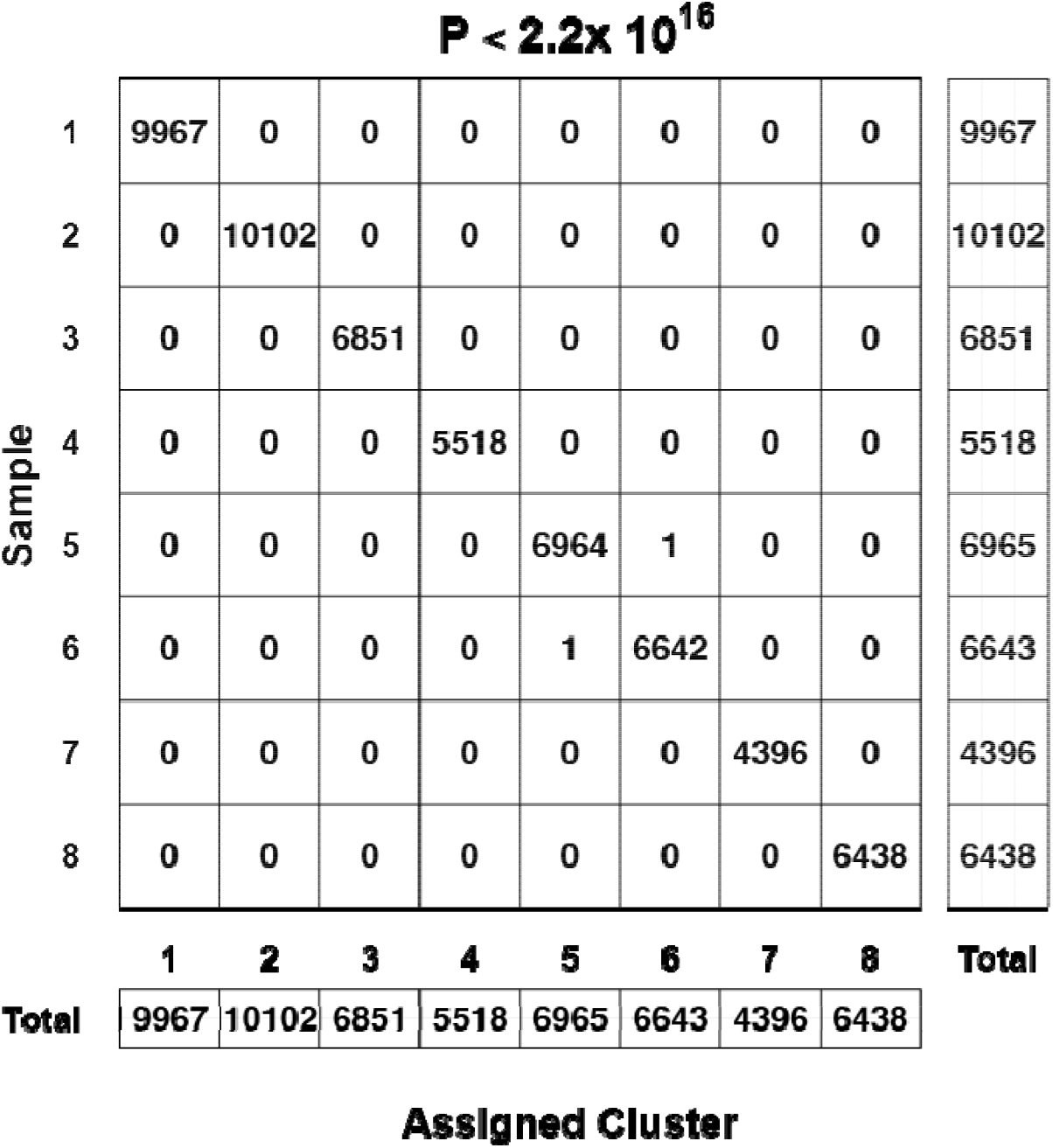
Inferred Cell Clusters are Highly Concordant with Expected Sample Identity. Contingency table showing numbers of cells from each independent donor sample (y-axis) and assigned to each inferred cluster (x-axis). Total cell numbers from each sample and assigned cluster shown in right and bottom table respectively. Independence of barcode-inferred cell identity and inferred cell cluster was tested *via* Pearson’s X^2^ test (p value depicted).

## Discussion

This simple workflow provides a straight-forward strategy for the deconvolution of sample identity from single-cell genomic data. Using genotype imputation to increase the availability of single-cell genetic information, followed by *k*-medoids clustering, we were able to recover donor identity for the vast majority of cells without use of barcoding data.

## Conclusion

These methods, utilizing natural genetic information from RNA-seq data for cell deconvolution, have the potential to provide rapid and enhanced quality-control for droplet-based barcoding. Indeed, if applied to barcoded data, this approach could aid in identifying sample cross-contamination or cell mis-labelling, whereby such cells would be clustered by their true donor of origin.

## Methods

### Genotyping, Phasing & Imputation

FASTQ files were aligned to the human genome reference sequence (hg38), before gene-level unique molecular identifier (UMI) quantification, as detailed(4). Post-alignment, cell barcodes with < 200 genes and < 1000 UMIs were filtered. Cells were then genotyped using cellsnp-lite v1.2.0(7) against a reference panel of 7.4 million SNPs identified through the 1,000 Genomes Project(5) (VAF >5%). Two samples had chromosomes for which no SNPs could be genotyped due to lack of read coverage, and these samples were excluded as a quality control measure. Default parameters were used, except that no filter on minimum inferred minor allele frequency was employed. Haplotype phasing and genotype imputation were then performed on a per-sample, per-chromosome basis with Eagle2 v2.4.1(8) and Minimac4 v1.0.2(9,10) respectively. Eagle2 was used with default parameters against a 1,000 Genomes reference panel as provided by the authors(11). (Minimac4 was run with the provided 1,000 Genomes (phase 3) M3VCF reference files(12) and with the --min-ratio parameter set to 10^−10^ to allow for regions with few genotyped SNPs. Imputed SNPs with an R^2^ value less than 0.9 were removed to increase accuracy and reduce downstream computational demand.

### Clustering

To cluster cells by genotype, a distance matrix was constructed by computing the pairwise Hamming distance between all cells from each sample. Hamming distance counts the number of positions that differ in two equal length strings, corresponding in our case to the number of genotypes which differ. Cells were then clustered into *k* clusters using the ClusteR R package v1.1.0(13) implementation of the Partitioning Around Medoids (PAM) algorithm(6). *k* was set to the number of input samples (eight). Cell cluster assignments were compared to cell sample of origin (inferred through sample-barcoding) to construct a contingency table, before assessing statistical independence of the two sources of cell labelling via Pearson’s X^2^ test. Data were visualized with BoutrosLab.plotting.general v7.0.3 in R v4.2.1.

## Declarations

### Ethics approval and consent to participate

The dataset used in this study was obtained from patients for whom informed consent was obtained previously(4).

## Availability of data and materials

The dataset supporting the conclusions of this article is available in the European Genome Archive, https://ega-archive.org/search/EGAD00001007677.

## Author contributions

R.G conducted all data analysis and implementation of the processing workflow. R.H provided guidance on data analysis and edited the manuscript. F.N prepared specimens for single-cell sequencing and contributed to initial data processing. P.C.B conceived of the workflow concept and provided guidance on implementation. All authors contributed to the final data interpretation and critical revision of the manuscript and approved the final version of the manuscript.

## Acknowledgments

Not applicable

## List of abbreviations

scRNA-seq: single-cell RNA-sequencing
dscRNA-seq: Droplet single-cell RNA-sequencing
PAM: partitioning around medoids
UMI: unique molecular identifier

